# A mathematical model of antibiotic resistance gene flow from livestock and spread amongst humans

**DOI:** 10.1101/2022.03.16.484526

**Authors:** Helen Fryer

**Affiliations:** Big Data Institute, University of Oxford, Old Road Campus, OX3 7LF

**Keywords:** Antibiotic resistance, bacterial infections, mathematical model, agriculture, livestock, animals, transmission, viral fitness, risk reduction, zoonotic, pathogens, epidemiology, foodborne

## Abstract

The evolution and spread of antibiotic resistance poses a major threat to human health. The high level of antibiotic use in the rearing of livestock is contributing to the origin and persistence of antibiotic resistance amongst humans. Understanding how resistance genes spread from livestock to humans and investigating the impact of managing antibiotic use in livestock will be important for guiding strategies to reduce the risk to humans. We have developed a mathematical model of the transmission of resistance genes from livestock to humans and their spread amongst the human population. Using this framework we demonstrate that although resistant zoonotic foodborne infections do not contribute significantly to the annual burden of death, they could be a source of resistance amongst other pathogens, including those that exclusively spread between humans. Amongst these pathogens, only livestock-derived resistant strains that are associated with a net fitness cost would be expected to decline in prevalence following control strategies aimed at reducing the impact on humans of antibiotic use in livestock.

**Author Summary:** Antibiotics are an essential component of human health care. They are also used in the rearing of livestock for human consumption, which contributes to the development of antibiotic resistant pathogens. Although humans are known to be at risk from antibiotic resistance that evolves amongst livestock, the number of human lives that are at stake remains unclear. Furthermore, the number of lives that could be saved through interventions to reduce antibiotic use in livestock has not been evaluated. Here, we have developed a mathematical framework to explore these questions. Using this framework we explicitly demonstrate that there are too many uncertainties to calculate the number of preventable human deaths. Nevertheless, the importance of reserving new and currently effective antibiotics for human use is clear. Once resistance genes that do not have a fitness disadvantage have spread from livestock to humans, it is too late for interventions targeted at livestock to affect their prevalence in humans in the long term.

## Introduction

The evolution and spread of antibiotic resistance presents a major and growing threat to the effective treatment of human bacterial infections and risks undermining many advances in human medicine (1, 2). Antibiotics are used in the rearing of livestock for human consumption, retrospectively to treat infections, prophylactically to maintain infection-free herds and for growth promotion. Their use is associated with the emergence and spread of antibiotic resistance, which has been identified in live farm animals (3-6), animal produce (7, 8) and environments contaminated with animal waste (9-12).

There is considerable overlap in the classes of antibiotics used in both livestock and humans (13), and there is a growing body evidence for the transfer of antibiotic resistance genes from livestock to human pathogens and to the human commensal population (14-16) (reviewed elsewhere (17, 18)). Small studies have revealed that resistance genes of Staphylococcus aureus transfer zoonotically between livestock and farmers (19, 20). Phylogenetic analysis of larger, population samples of human and animal staphylococcus aureus has also provided convincing population evolutionary evidence of the transmission of resistance from animals to humans (21, 22). Indistinguishable vancomycin-resistant enterococci (VRE) has been identified in the faeces of a Dutch farmer and his turkey flock (23); and on a wider scale, extensive use of vancomycin as a growth promoter in livestock rearing in Europe has been associated with faecal carriage of VRE in healthy humans (24).

Humans are at risk of acquiring antibiotic resistance genes from livestock through different paths. Humans can be directly infected with resistant pathogens (notably Salmonella (25), Campylobacter (26) and *e-coli* (27)) as a result of the consumption of infected meat or other food products that have become contaminated through the environment. However, resistance genes can also transfer ‘horizontally’ (28-30) between different strains of the same bacterial species (31, 32), and in some cases, different bacterial species (32, 33). As a result of gene transfer, even pathogens that are spread entirely amongst humans are at risk of acquiring resistance genes from animal pathogens following sporadic zoonotic infection from a food source or through close animal contact during farming. This could result from the co-infection of an individual with both a human and animal pathogen. Alternatively, an animal pathogen could transfer resistance genes into an individuals’ commensals (notably enterococci, Staphylococcus aureus or *e-coli*). Once in the commensals, resistance genes can linger for long periods (34), over which there remains a risk of incorporation into an infecting human bacterial pathogen of similar or different species (35). The gut is an optimal site for gene transfer, especially to human pathogens that infect the gastrointestinal tract, because bacterial populations there are large in number and share a similar ecology (36, 37). Because horizontal gene transfer allows the exchange of combinations of mutations, it can potentially facilitate the spread to different pathogens of highly adapted resistance genes that may have taken a long time to evolve in one pathogen. The evolution and spread of an extensively drug-resistant clone of the exclusively human enteric infection, Salmonella Typhi, has recently been shown to have resulted from the acquisition of a plasmid harbouring genes resistant to fluoroquinolones and cephalosporins (38). Although the study did not explore whether these resistance genes derived from the use of antibiotocs in humans or livestock, both antibiotic classes have been used in agriculture and have been associated with antibiotic resistance in food-producing animals (39, 40). This finding highlights the potential risk that the use of antibiotics in livestock poses to human health.

It has been argued that reducing the use of antibiotics in livestock will reduce the introduction and spread of antibiotic resistance amongst humans (41, 42). Sweden led the way in implementing risk reduction strategies and discontinued the use of antibiotics as growth-promoters in 1986 (43). In 1999, similar restrictions were rolled out across the EU (44). Often the impact of local policy is unquantified; however, surveys that have been published reveal limited direct benefits of interventions on human health (45). For example, following the 1995 ban on vancomycin as a growth promotor in Germany, a reduction from 12% in 1994 to 3% in 1997 in the faecal carriage of vancomycin-resistant enterococci in humans was recorded (24), but this was not reflected in a reduction in pathogenic human infections with VRE (24). An explanation for this is that when vancomycin-resistant enterococci infections persist in humans, they can often be linked to human use of antibiotics (46) and transmission between humans in a hospital setting (47). In the United States of America (USA), no decline in the levels of ciprofloxacin-resistant Campylobacter in chickens (48, 49) or humans (CDC. National Antimicrobial Resistance Monitoring System (NARMS) Now: www.cdc.gov/narmsnow) has been observed following the 2005 ban of fluoroquinolones in chickens. It has been argued that it this is due to the absence of any fitness cost to resistance (50).

These examples highlight that it is not straightforward to predict how reducing the spread of resistance genes from livestock to humans will change the prevalence of resistance in humans. In particular, the impact will be different for different pathogens and for different resistance genes. Here we develop a mathematical model to explicitly characterize the different ways in which antibiotic resistance genes can spread from livestock to humans (51, 52). We use the model to investigate the circumstances under which a reduction in antibiotic resistant infections in humans could be achieved by reducing the spread of antibiotic resistance genes from livestock to humans. We investigate how the following factors change the impact on humans of restricting gene flow from livestock to humans: the transmission dynamics of the pathogen (for example, whether the pathogen is transmitted to humans mostly from livestock or mostly from other humans); how resistance is acquired; the fitness profile of resistance; and the relative rate that resistance is acquired from livestock and alternative (i.e. human) sources. The timing of interventions, that is, whether they are applied before or after resistant strains have spread to humans, is also considered.

## Results

### A new mathematical model of antibiotic resistance gene flow from livestock and spread amongst humans

We developed a new mathematical model of the transmission of antibiotic resistant genes from livestock to humans and their spread amongst a human population. The model focusses on the dynamics of resistance associated with a single (unspecified) bacterial pathogen and there is competition between sensitive and resistant strains of the pathogen. The model can be parametrized to consider pathogens with different dynamics in regards to how the pathogen is transmitted and how resistance genes are acquired from livestock. One example is a pathogen that is primarily transmitted to humans from livestock and resistance can be acquired directly through transmission of a resistant pathogen (Fig 1a). A second example is a pathogen for which the majority of infections derive from humans, but a minority derive from livestock and for which resistance can be acquired through transmission of a resistant pathogen (Fig 1b). A third example is a pathogen that exclusively spreads to humans from other humans and for which resistance genes, deriving from an animal pathogen of a similar or disparate genus, can be acquired via gene transfer following coinfection, or via a commensal route (Fig 1c). In the model, resistance can also derive from two non-livestock-related sources: *de novo* mutation and gene transfer from an alternative source, for example, a different human pathogen.

**Fig 1.**
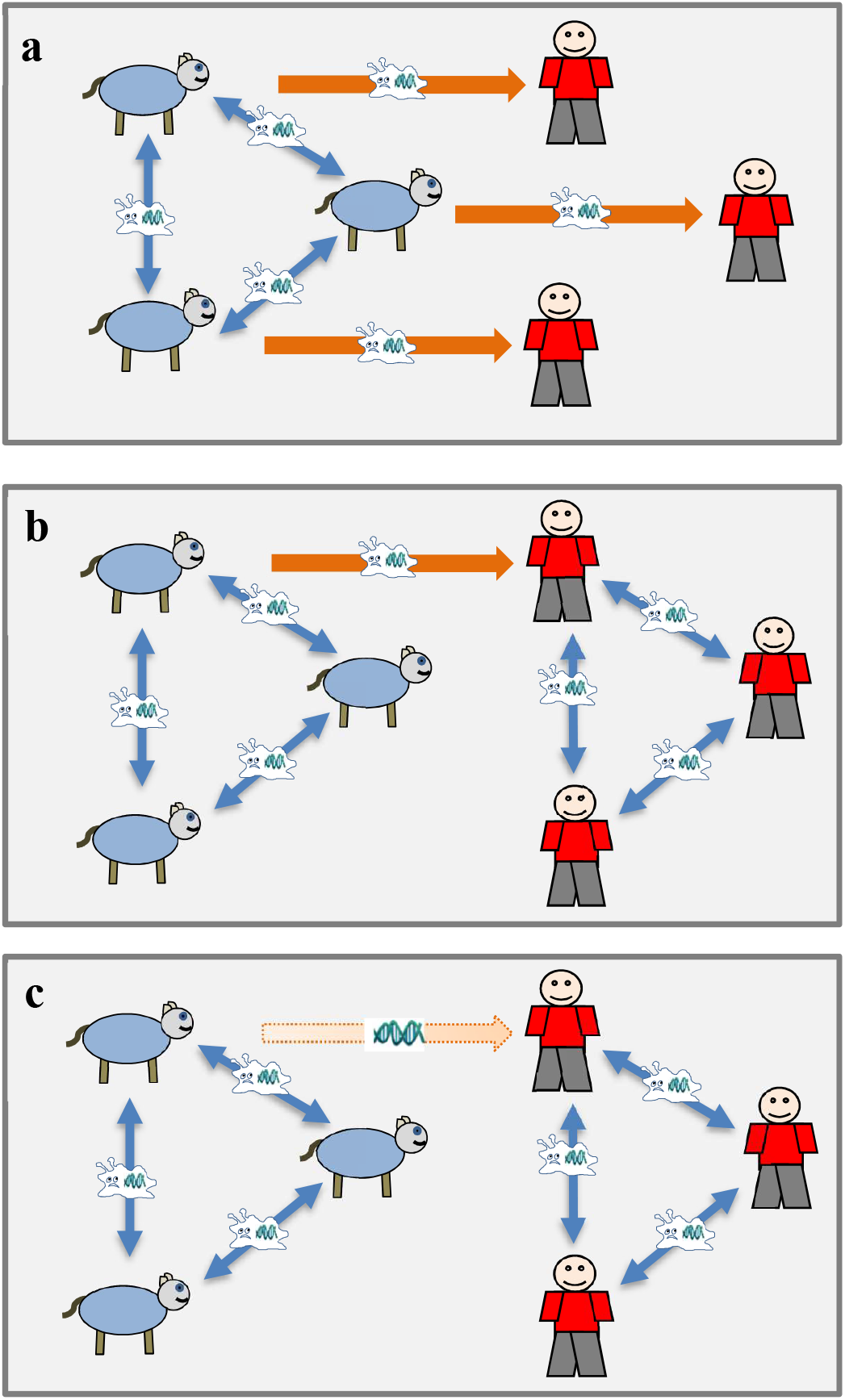
There is variability between human bacterial pathogens in how humans acquire the pathogen and how resistance is acquired from animals. This figure highlights three example pathogens, categorized in terms of how humans are impacted by antibiotic resistance in livestock. In a) the majority of human infections result from direct transmission from animals (e.g. a foodborne infection) and the antibiotic resistant pathogen is transmitted directly from animals to humans. In b) the majority of human infections result from transmission of the pathogen from other infected humans. A minority result from direct transmission from animals, during which the antibiotic-resistant pathogen may be transmitted. Once the resistant pathogen is in humans, it can be transmitted between humans. In c) all human infections result from transmission from other infected humans, however, resistance genes can transfer from livestock to humans leading to a resistant pathogen in humans that can be transmitted to other humans.

The model explicitly acknowledges that some resistance mutations impose a fitness cost on the pathogen (53-55) whereas others have greater stability in the absence of antibiotic use (56, 57). Fitness costs are known to impact the dynamics of resistant pathogens in one or more ways; for example, reduced transmissibility to untreated individuals, a reduced infectious period in untreated individuals or reversion to a sensitive strain following transmission to an untreated individual. However, in each case, increased fitness costs favour the existence of sensitive, rather than resistant strains in the absence of antibiotic treatment. We model a fitness cost by allowing for the reversion of resistance amongst untreated individuals.

Antibiotic resistance may also be associated with a fitness advantage amongst antibiotic-treated individuals in the form of slower recovery rates or increased transmissibility. Our model allows for such a fitness advantage through different rates of recovery. The model is illustrated in Fig 2 and presented in detail in the Methods section. Model variables and parameters are listed in Table 1. We use our model to investigate how resistance amongst livestock can contribute to the emergence and spread of resistant infections in humans. We also investigate the circumstances under which reducing the spread of resistance genes from livestock to humans (for example through reducing antibiotic use in livestock) can reduce resistant infections amongst humans.

**Table 1.** Model variables and parameters.

| Variables |  |
| --- | --- |
| $t$ | Time |
| $U(t)$ | Number of untreated individuals who are infected with a drug-sensitive strain |
| $\tilde{U}(t)$ | Number of untreated individuals who are infected with a drug-resistant strain |
| $T(t)$ | Number of treated individuals who are infected with a drug-sensitive strain |
| $\tilde{T}(t)$ | Number of treated individuals who are infected with a drug-resistant strain |
| $S(t)$ | Number of individuals who are susceptible. |
| Parameters |  |
| $\pi$ | Fraction of individuals who receive treatment immediately upon infection |
| $\Lambda$ | The per-person rate that susceptible individuals become infected with a pathogen directly from an infection in livestock |
| $\alpha$ | The fraction of human infections with an animal pathogen that are resistant |
| $\beta$ | Transmission coefficient. This encompasses the degree of contact between susceptible and infected individuals and the probability of infection per contact. |
| $\mu$ | The per-person rate of *de novo* emergence of drug resistance in treated infected individuals |
| $\phi_N$ | The per-person rate that treated infected individuals acquire resistance through gene transfer from a non-livestock source. |
| $\phi_A$ | The per-person rate that treated infected individuals acquire resistance through gene transfer from livestock |
| $\psi$ | The per-person rate of reversion of drug resistance in untreated individuals |
| $\delta$ | The per-person rate of recovery of untreated infected individuals |
| $\sigma$ | The per-person rate of recovery of treated individuals infected with a drug-sensitive strain |
| $\varepsilon$ | The per-person rate of recovery of treated individuals infected with a drug-resistant strain. |
| $z_1$ | The effectiveness of strategy 1 |
| $z_2$ | The effectiveness of strategy 2 |
| $N$ | Total number of individuals in the population |

**Fig 2.**
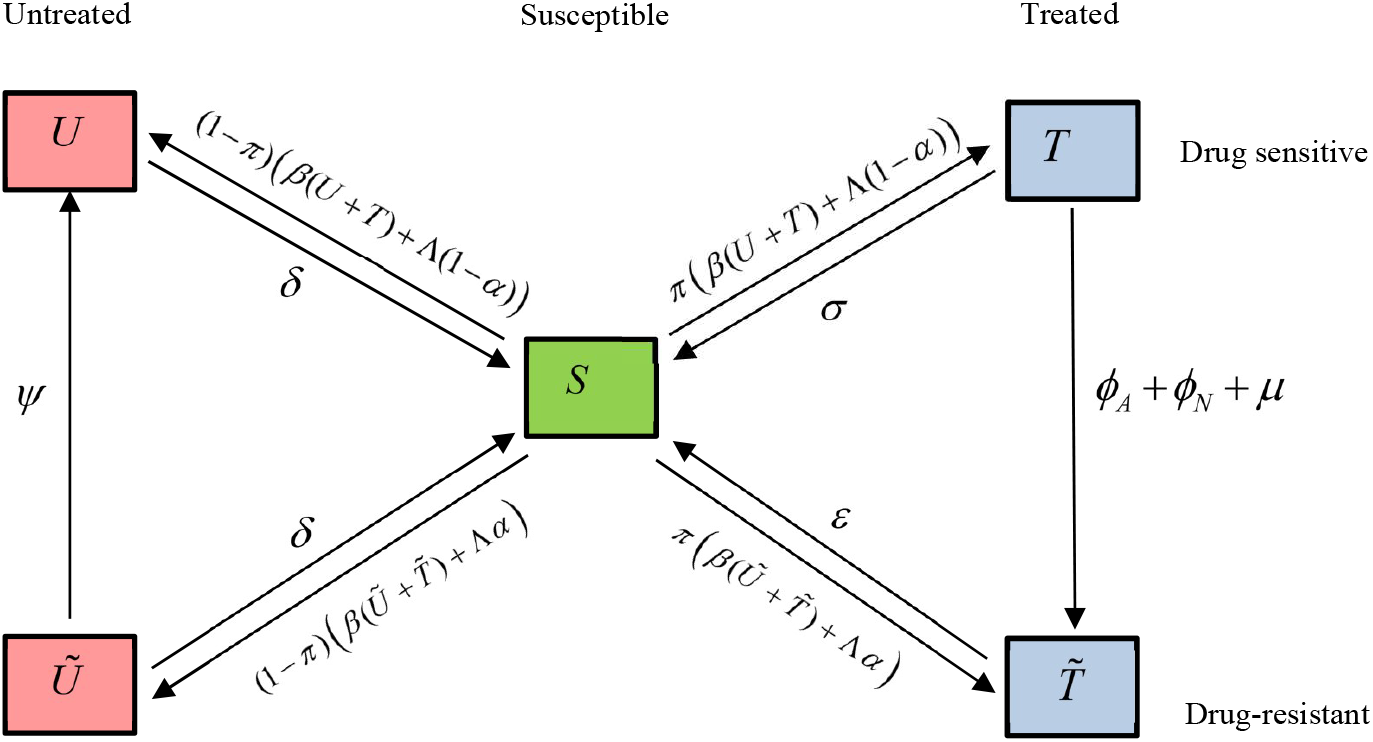
Model 1. A mathematical model of antibiotic resistance gene flow from livestock and its spread amongst humans. This model tracks the spread of a bacterial pathogen amongst a human population, a portion of which receive antibiotic treatment upon infection. Infections can result from transmission from other infected humans or from exposure to infections in livestock. There is competition between an antibiotic-sensitive and an antibiotic-resistant strain. Resistant strains can emerge in treated individuals via de novo mutation or gene transfer deriving from livestock or from an alternative (i.e. human) source. In addition, resistant pathogens can be transmitted to humans from animals directly. Resistant strains can revert in untreated individuals. Recovery of infection occurs at different rates according to the pathogen strain and the treatment status. The infection rates shown here assume that there is no intervention to reduce the spread of resistance from animals to humans although the full model includes the impact of interventions (see Methods).

### Antibiotic resistance in livestock can contribute to the emergence and spread of resistance amongst human bacterial infections, even for pathogens that exclusively spread amongst humans

To understand how antibiotic resistance in livestock can contribute to resistance in humans, we begin by parametrizing the model to represent a pathogen for which all human infections are acquired from other humans (Fig 1c). Fig 3 shows simulated epidemics in which, prior to time 0, the infection has reached equilibrium amongst humans and antibiotic resistance is absent. At time 0, a resistant strain can be acquired through three different routes: *de novo* mutation in treated humans and horizontal transfer of resistance genes that derive from livestock or horizontal transfer of resistance genes from an alternative (i.e. human) source. We compare different scenarios in which the relative magnitudes of acquisition of resistance from livestock, versus an alternative source differ and in which the fitness profile of the resistant strains differ. These simulations show that even for pathogens that are spread entirely amongst humans, resistance in livestock can contribute to the evolution of resistance amongst humans via gene transfer from animal pathogens, for example, foodborne pathogens (compare red and blue solid lines, in each panel). They also highlight that the fitness profile of the resistant strain impacts the spread of resistance. For resistant strains that confer a fitness advantage in treated individuals or do not confer a fitness cost in untreated individuals (Fig 3a, 3b and 3d), resistance in livestock speeds up the spread of resistance in humans (to fixation if there is no fitness cost (Fig 3a-b) or to an equilibrium prevalence less than 100% otherwise (Fig 3d)). For resistant strains that are associated with a negligible fitness advantage in treated individuals and a measurable fitness cost in untreated individuals, resistance in livestock not only speeds up the spread of resistance, but it also leads to a higher equilibrium prevalence of resistance amongst humans (Fig 3c). In each context, however, resistance in livestock only measurably contributes to the spread of resistance amongst humans provided the rate that it is acquired is comparable to the rate at which it is acquired from alternative sources (e.g. via *de novo* mutation or gene transfer from another human pathogen).

**Fig 3.**
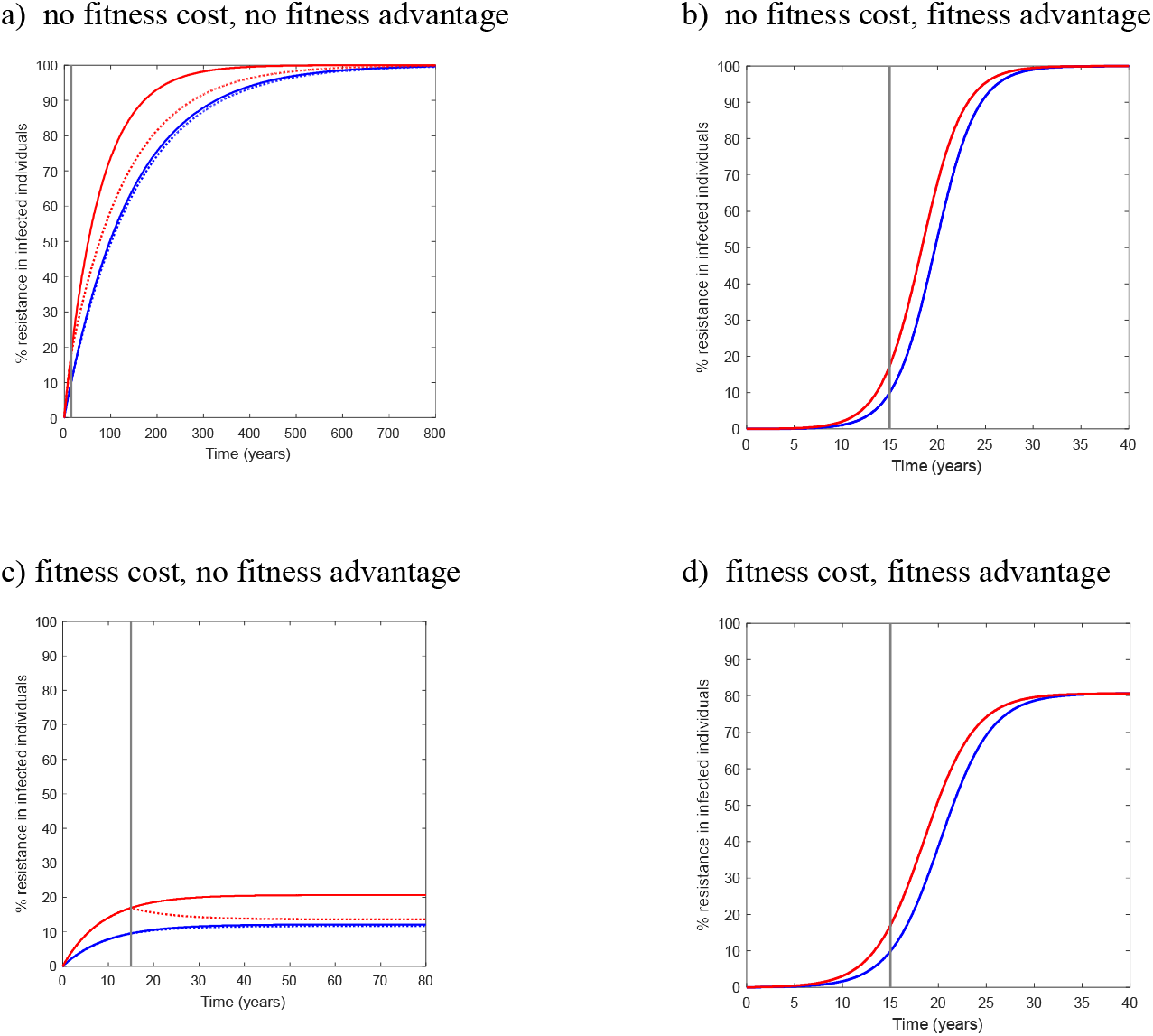
Model predictions of the impact of the transfer of resistance genes from livestock to humans on the spread of resistance amongst an exclusively human-human spread pathogen. This figure illustrates how the transfer of resistance genes from livestock to humans can contribute to the spread of resistance among humans. It also highlights how, once a resistance gene begins to circulate amongst humans, interventions that reduce the prevalence of resistance in livestock will only have a long-term impact upon resistance prevalence amongst humans provided there is a net fitness cost to resistance. Across panels a)-d) different scenarios in regards to the fitness profile of the resistant strain are compared. The fitness cost of resistance in the absence of treatment is compared by varying the reversion rate. In a) and b) there is a no fitness cost (Ψ = 0 years^-1^) and in c) and d) there is a measurable fitness cost (Ψ = 0.1 years^-1^) in untreated individuals. The fitness advantage of resistance in the presence of treatment is compared by varying the recovery rate of treated resistant infections. In a) and c) there is a no fitness advantage ε = 24 years^-1^), and in b) and d) there is a measurable fitness advantage (ε = 20 years^-1^), in treated individuals. In each panel, two rates at which the resistance gene spreads from livestock to humans are compared (solid lines) to represent differences in the prevalence of resistance in livestock. In blue, the ratio of the rate that resistance is acquired from livestock versus an alternative source (ϕ_A_ : (ϕ_N_ + μ)) is 5:95, i.e. 5% of resistance originates from livestock and 95% from an alternative source. The rates of ϕ_A_ and ϕ_N_ + μ differs between a, b, c and d to yield a resistance prevalence of 10% at year 15 when 5% of resistance derives from livestock. In red, the rate ϕ_A_ is increased by a factor of 19, but the rate ϕ_N_ + μ remains unchanged, so that the ratio of the rates is 50:50. In each figure, at year 15 (grey line), an intervention to reduce the rate that resistance is acquired from livestock by gene transfer by 80% (z_1_ = 0.8) is applied (these appear as dashed lines in a and c, but are equal to the solid lines and therefore not visible in b and d). In a) eventual fixation of the resistant strain is inevitable. Reducing the rate that resistance is acquired from livestock can delay the spread of resistance, provided the spread of resistance from livestock contributes measurably to the generation of resistance in humans. In b) eventual fixation of the resistant strain is inevitable. Reducing the rate that resistance is acquired from livestock has no impact upon the prevalence of existing resistant strains. In c) the resistant strain does not reach fixation. Reducing the rate that resistance is acquired from livestock can measurably reduce the prevalence of resistance in humans provided that resistance in livestock contributes significantly to the generation of resistance in humans (compare solid and dashed red lines). In d) the resistant strain does not reach fixation. Reducing the rate that resistance is acquired from livestock has no measurable impact upon the prevalence of existing strains. The following parameters are used in each simulation: β = 14.26, π = 0.3, σ = 24 years^-1^, δ =12 years^-1^ and Λ = 0 years^-1^. Furthermore, in a), b), c) and d) respectively, μ +ϕ N = 3.6 ×10^−3^, 4.1×10^−4^ and 0.122 years^-1^ and in blue ϕ A = 3.6 ×10^−3^, 2.18 ×10^−5^, 6.4 ×10^−3^ years^-1^ and 6.5×10^−5^. Results are independent of N (i.e. choose any N).

For pathogens for which the majority of infections are transmitted to humans from other humans and the minority are transmitted from animals (Fig 1b), the dynamics of resistance are similar (Fig S1). However, the eventual fixation of resistant strains that do not confer a fitness cost in untreated individuals is not inevitable. If there is no net fitness advantage to resistance, resistance derived from livestock can change the equilibrium prevalence of resistance (Fig S1a).

For pathogens for which the majority of infections are transmitted to humans from animals (Fig 1a; i.e. zoonotic foodborne pathogens, such as Campylobacter or Salmonella) resistance prevalence in humans does not gradually build up or decline over time; instead, it reacts directly to the factors that dictate the prevalence. Assuming that most resistance is transmitted directly at infection, the prevalence of resistance in humans is principally governed by the prevalence of resistance amongst animals and the fitness costs of resistance. However, it is noteworthy that some resistant zoonotic foodborne human infections may be the result of an infection with a sensitive foodborne pathogen, followed by the within-host emergence of a resistant strain (via *de novo* mutation or gene transfer of resistance that derived from a different human or animal pathogen). In such a circumstance, the resistance prevalence amongst humans is dependent upon the rate of resistance emergence within the human host.

### For pathogens that exclusively spread amongst humans, once a resistant gene that has no net fitness cost has emerged within a human pathogen, its eventual fixation cannot be prevented by reducing the spread of that gene from animals to humans

The fact that antibiotic resistance derived from livestock contributes to the spread of resistance amongst humans points to the need for interventions to manage this risk. The expected impact of such interventions, however, is not straightforward; rather it depends upon context. For example, for pathogens that are spread to humans entirely from other humans, interventions to reduce the prevalence of resistance in livestock cannot prevent the eventual fixation of resistant strains that have already been transmitted to humans and which are associated with no fitness cost (Fig 3a and 3b dashed lines). At best, such interventions could delay fixation (compare dashed and solid red lines in Fig 3a). Only for resistant strains that are exclusively generated in livestock could the introduction and spread to fixation be completely prevented through interventions targeted at livestock. In such a scenario, it would be necessary to entirely stop the resistance gene from spreading from animals to humans.

### Reducing the spread of antibiotic resistance from animals to humans can lead to a reduction in the number of human resistant infections in certain contexts

It is intuitively clear that the impact of interventions to reduce the spread of antibiotic resistance from livestock to humans depends upon a range of factors. For example, it depends upon the transmission dynamics of the pathogen, whether the resistance gene already prevails amongst the human pathogen, the fitness profile of resistance and the relative degree to which resistance amongst humans derives from livestock and alternative sources. Here, we explore in more detail how the expected impact of interventions that are targeted at livestock on human infections with an existing antibiotic-resistant strain is dependent upon such factors.

To achieve this, we adapted the mathematical model to track the number of infections for which resistance derives from livestock versus an alternative (i.e. human) source. This adaptation (Fig S2) was made because rate estimates for the various processes that contribute to the generation of resistance are not typically available. However, genetic analysis and phylogenetic reconstruction of sampled bacterial strains offer a potential means to investigate whether or not a strain derives from livestock. Estimates of the proportion of sampled resistance – amongst a particular human pathogen – that derives from livestock is, therefore, an observation that could plausibly be made and used for understanding the impact of interventions.

Fig 4 shows model predictions of the expected impact of interventions on the number of human resistant infections under different parametrizations of the model that dictate the transmission dynamics of the pathogen, the fitness profile of resistance and the proportion of observed resistant infections that derive from livestock. In each simulation used to generate these results, prior to time 0, an infection has reached an equilibrium prevalence of 1% amongst humans and antibiotic resistance is absent. At time 0, antibiotic resistance emerges amongst humans through livestock and alternative routes at such a (combined) rate that 15 years later, 10% of infections are resistant.

**Fig 4.**
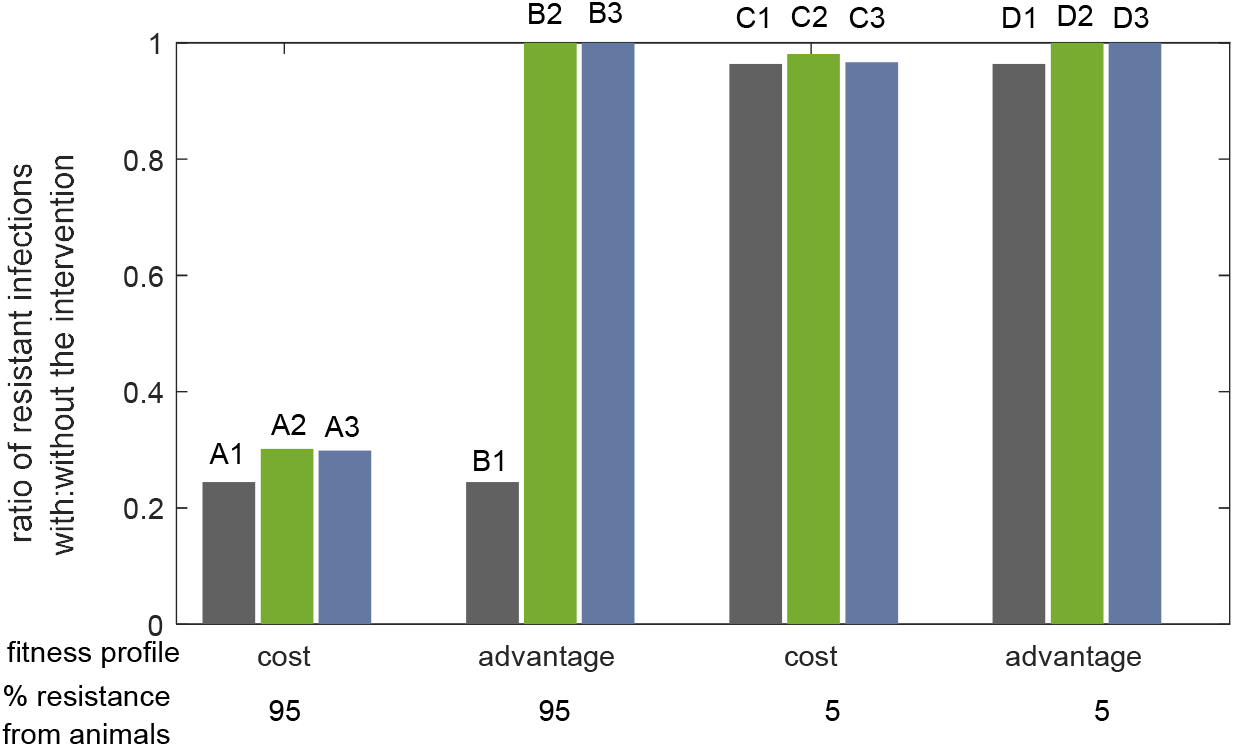
Whether the number of human infections with an existing antibiotic resistance strain can be reduced by reducing the prevalence of resistance in livestock depends upon different factors. This figure explores the impact that reducing the prevalence of resistance amongst animals (intervention 1) by 80% would have on the number of human infections with an existing antibiotic resistant strain, according to different factors. These factors are the transmission dynamics of the pathogen, the fitness profile of the resistance strain and the percentage of observed resistance that derives from livestock. The coloured bars categorize whether the pathogen is transmitted to humans primarily (99.9%) from animals (grey), primarily (99.9%) from humans (green) or entirely from humans (blue). The fitness profile is varied to consider a resistant strain that either has a net fitness cost to resistance (measurable fitness cost in untreated individuals and negligible fitness advantage in treated individuals) or a net fitness advantage to resistance (negligible fitness cost, measurable fitness advantage). In each model simulation used to generate the results shown, the model is independently parametrized to ensure that prior to time 0, an infection has reached a prevalence of 1% amongst humans and antibiotic resistance is absent. At time 0, antibiotic resistance emerges amongst humans at such a rate that 15 years later, 10% of infections are resistant. In one option, 95% of resistance observed at year 15 originates from livestock, whereas in the other option the value is 5%. The coloured bars show the ratio of the number of resistant infections at year 50 with:without the intervention. The parameters used in these simulations are listed in Supplementary Table 1.

An intervention (intervention 1) is then modelled in which at year 15, the prevalence of resistance amongst animals is reduced by 80%, although the total prevalence of bacterial infections that are transmitted to humans does not change. This could represent a reduction in antibiotic use amongst animals and the deployment of alternative strategies to manage bacterial infections.

Figure 4 shows the impact of this intervention on the number of resistant human infections at year 50. It highlights that for pathogens that transmit to humans primarily (green bars) or entirely (blue bars) from other humans, if resistant strains confer a net fitness advantage (e.g. a negligible fitness cost in untreated individuals and a measurable fitness advantage in treated individuals), the intervention yields no reduction in the number of resistant infections by either measure (scenario B2 and B3).

By contrast, if there is a measurable net fitness cost to resistance (e.g. negligible fitness advantage in treated individuals and a measurable fitness cost in untreated individuals), a reduction in the number of resistant human infections is predicted (scenarios A2, A3, C2 and C3). The expected reduction is greater when the portion of resistance that derives from an animal source is greater (compare scenario A2 with C2 and compare scenario A3 with C3).

For pathogens that are transmitted to humans primarily from animals (i.e. zoonotic foodborne pathogens) (grey bars in Fig 4) interventions that are aimed at reducing resistance in livestock can reduce the number of resistant human infections, irrespective of the fitness profile. Once again, the reduction is larger when the proportion of resistance that derives from livestock is larger (compare scenario A1 ≈ B1 with scenario C1 ≈ D1). Table 2 summarises these findings.

**Table 2.** What is the expected impact of reducing the spread of antibiotic resistance from livestock to humans upon the number of human infections with an existing resistance strain? This table categorizes the impact of intervention 1 upon the number of resistant human infections according to different factors. *The magnitude of the impact is larger if the proportion of resistance that derives from livestock is larger (see Fig S4 for details).

| Is there<br>a net fitness<br>cost or advantage<br>to resistance? | How is the pathogen transmitted to humans? |  |  |
| --- | --- | --- | --- |
|  | Majority from<br>animals | Majority from<br>humans | All from<br>humans |
| Cost | Reduce resistant<br>infections* | Reduce resistant infections* |  |
| Advantage |  | No significant impact |  |

The impact of this intervention on resistant strains that have a neutral fitness profile (e.g. no fitness cost and no fitness advantage) as well as the impact of a second intervention (intervention 2) in which the total number of human resistant infections that derive from animals is reduced is explored in Fig S3. Figure S3 also explores the long term impact (i.e. at equilibrium). The magnitude of the reduction in the equilibrium number of human infections that results from each intervention is explored in Fig S4.

### Towards a quantitative estimate of the impact on human health of reducing the spread of antibiotic genes from livestock to humans

Policymakers are interested in evaluating the number of infections and deaths due to resistance that could be prevented by reducing the spread of antibiotic resistance genes from livestock to humans. To this end, it is useful to consider how different pathogens contribute to the total burden of existing antibiotic-resistant infections amongst humans and which of these infections could be prevented through interventions targeted at livestock. It is also crucial to consider the role that different pathogens play in the generation and spread of new resistance genes to bacterial pathogens. In Table 3 we present a list of important resistant human bacterial pathogens in the United States. For each pathogen, we reproduce published data of the minimum estimated numbers of human infections (Table 3, column 3) and deaths (Table 3, column 4) due to drug-resistance in the United States in 2010 (58). We note that estimates of the global burden of resistance are called for, but that in the absence of such data the data for the US are useful for illustrating our concept.

**Table 3.**
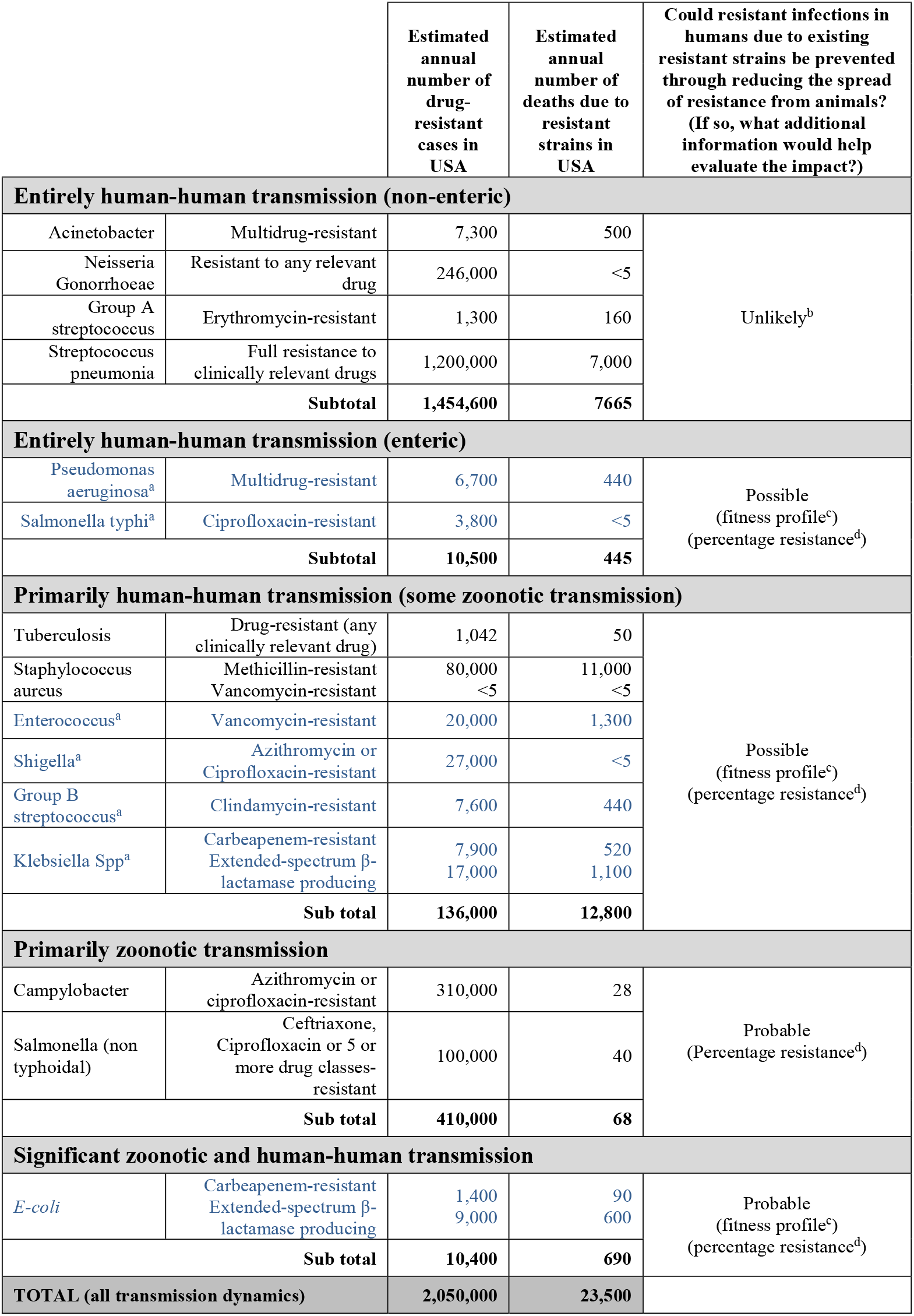
Antibiotic resistance in important bacterial pathogens and information to aid risk reduction. This table lists minimum annual estimates of human morbidity and mortality from antibiotic-resistant infections in important human bacterial pathogens in the United States in 2013 (58). Pathogens are classified according to how the pathogen is transmitted to humans and whether it is enteric. Information required to estimate the impact on humans of livestock-targeted interventions are listed. ^a^Enteric infections are shown in blue. These infections are at a greater risk than non-enteric infections of acquiring resistance genes from livestock through gene transfer as a result of foodborne infections. ^b^It is unlikely, although not impossible that a non-enteric pathogen that is entirely transmitted between humans can acquire resistance from an animal pathogen. Gene transfer following coinfection with an animal pathogen that populates a similar anatomical niche is possible. ^c^The fitness cost of resistance in untreated individuals and the fitness advantage of resistance in treated individuals. ^d^The percentage of observed human resistant infections for which resistance derives from livestock.

Our study highlights that understanding the transmission dynamics of different human pathogens can contribute to understanding how interventions to reduce the spread of antibiotic resistance genes from livestock to humans can affect resistance prevalence in humans. For most pathogens this information is reasonably well characterised, at least in terms of whether or not human infections are 1) mostly acquired from other humans, 2) entirely acquired from other humans or 3) mostly acquired directly from animals as foodborne infections. To demonstrate how far we can get in addressing the potential impact of reducing the spread of antibiotic resistance genes from livestock to humans, the important human pathogens listed in Table 3 are classified according to their transmission dynamics. We also highlight (in blue) those which are enteric as these pathogens are more likely to exchange genes with zoonotic foodborne pathogens than non-enteric pathogens. Our study also highlights that other information is typically necessary to understand the likely impact of interventions and such necessary additional data for each of the listed pathogens is highlighted in column 5.

This representation highlights that pathogens that transmit to humans entirely from other humans and do not typically infect the gastrointestinal tract are responsible for the majority of human resistant infections (71%, n=1,454,600) and a significant proportion of deaths due to resistant infections (32%, n=7670). It is unlikely, though not impossible, that resistance from livestock contributes to resistance in these pathogens and therefore unlikely that interventions aimed at livestock could reduce resistant infections in these pathogens.

Enteric pathogens that transmit to humans entirely from other humans and pathogens that are primarily spread to humans by other humans, but sometimes spread by animals are responsible for only 7% (n= 146,500) of resistant infections but 56% of deaths due resistant infections (n=13,250). Our model reveals that reducing the spread of resistance from livestock to humans is particularly important for delaying the introduction of fit resistant strains to pathogens in these categories. However, to understand whether interventions could have an impact on existing resistant strains in humans requires additional data. Firstly, the fitness profile of the strain. Secondly, for those with a net fitness cost to resistance, what fraction of resistance derives from livestock. We would also need to know which animal pathogen is contributing to the human resistance. Currently, this range of information is typically unavailable, although genetic studies could be employed to gather it.

Pathogens that are principally transmitted to humans from animals, with little ongoing human-to-human transmission are responsible for a minority of infections (20%; n=410,000) and deaths due to resistance (0.3%; n=68). For these pathogens, resistant infections that are the result of direct infection of a resistant strain of the infecting pathogen could be reduced in number by reducing resistance prevalence in livestock. It is noteworthy, however, that even for these pathogens, some resistance in humans may derive, post infection, from an alternative source to livestock; notably *de novo* mutation following treatment, or gene transfer of resistance that derived from a human source. Interventions would only reduce these (post-infection-derived) resistant human infections if they reduce the total number of humans infected with the pathogen (refer to intervention 2 in Fig S3 and Fig S4). Although resistant pathogens in this category are responsible for only a minority of human deaths due to resistance, their potential role in the transfer of resistance genes to primarily (or entirely) human-human transmitted pathogens, as a result of co-infection, or via the commensals must not be overlooked.

*E-coli* is classified in a transmission category of its own because although it is primarily transmitted to humans from animals, either through food (52%) or direct contact (3%), other transmission routes are significant (14% human-human, 9% waterborne; 21% of infections have unknown transmission routes) (59). It is not clear what fraction of resistant infections are the result of direct infection with resistant *e-coli* from livestock – the portion that are could be directly prevented by reducing the spread of antibiotic-resistant *e-coli* from livestock to humans. For the remaining portion, more information, including the fitness profile of resistance and the fraction of observed resistance that is due to animals would be required to understand the impact of interventions through modelling.

In summary, more data are required to quantitate the impact that reducing the spread of antibiotic resistance genes from livestock to humans could have on infections and deaths due to existing antibiotic-resistant pathogens. Nevertheless, reducing the spread of antibiotic resistance genes from livestock to humans is universally beneficial in terms of delaying the introduction of new resistant strains to humans, in particular, those that are not associated with a net fitness cost.

## Discussion

In this study, we have used a mathematical model to show that even for pathogens that are spread entirely between humans, resistance derived from livestock can contribute to resistance amongst humans. Although zoonotic foodborne infections are directly responsible for only a minority of deaths due to resistance, reducing these infections could nevertheless delay the emergence of resistance genes amongst principally human-human spread bacterial pathogens. This is because foodborne infections provides a point of entry for resistance genes to enter the human gut where there exists a risk of their horizontal transfer to other human bacterial pathogens, particularly enteric ones. Delaying the introduction of resistance genes that are not associated with a net fitness cost is particularly important because once they have started to spread amongst pathogens that transmit between humans, interventions cannot reduce their prevalence in the long-term.

Reducing the spread of antibiotic resistance genes from livestock to humans can measurably reduce the prevalence of existing resistance strains amongst humans under particular conditions. For pathogens that transmit to humans primarily from livestock, it is sufficient that the fraction of resistance that derives from livestock is significant. For pathogens that transmit to humans primarily from other humans, if there is additionally a net fitness cost to resistance, the expected reduction is also measurable.

We intentionally do not model the within-host evolution and between-host spread of antibiotic resistance amongst livestock. Instead, we simply model the infection of humans with resistant animal pathogens and the transfer of resistant genes from animals to humans as rate constants. This generalisation acknowledges that there are different means of reducing the spread of resistance genes from livestock to humans. One means is to reduce the prevalence of antibiotic resistance amongst livestock through reduced antibiotic use. However, only resistance that is associated with a net fitness cost in livestock will decline in prevalence amongst livestock as a result of such an intervention. What is more, if applied in isolation, interventions to reduce antibiotic use could increase the total prevalence of infection amongst livestock and thereby increase infection rates amongst humans (12). Another means of reducing the spread of resistance genes to humans is to improve bacterial testing of food during processing or to reduce the overall burden of infection amongst livestock (60). The latter of these could be achieved through improved hygiene control (on farms, during livestock transport and in food preparation), vaccination, and rapid testing followed by culling (61). Vaccines currently exist for most of the principal diseases of livestock (62), excluding those which have a near-commensal association with their host species. However, costs are reportedly impeding vaccination uptake (60) and the commercialisation of new vaccines (63). Prebiotics, probiotics (64) and phage therapy (60, 65) are promising alternatives, but further development of these agents and demonstration of their economic and scientific viability is needed.

Consumer demand is likely to form part of the economic calculations determining whether to reduce antibiotic use in livestock. Consumer demand for reduced antibiotic use in food production is gaining support. Part of this support relates to concern over the spread of antibiotic resistance; however, arguably a more significant element in this growing movement is the desire for food that is perceived as naturally produced and free from residues (66).

By demonstrating that reducing the spread of resistance from livestock to humans can differentially impact the dynamics of resistance to different human pathogens, we highlight that empirical outcomes of antibiotic regulation of one pathogen should not be taken as proof-of-principal of the impact of regulation upon all. As we have highlighted, multiple factors that are specific to each resistant pathogen should be considered in the development of policy guidance, particularly in regards to existing resistant strains. More data are clearly required to make effective decisions. These include the fitness profile of resistant strains and an understanding of the contribution that livestock plays in driving resistance, relative to alternative sources.

Our model focusses on how within-host evolution and between-host transmission affects the prevalence of infection and antibiotic resistance, and therefore purposely does not explicitly measure deaths due to infection. However, policy guidance must also consider the degree of morbidity and mortality associated with the resistant and sensitive strains of each pathogen. The importance and irreplaceability of the antibiotics used to treat each pathogen is also crucial. Antibiotics of critical importance for human health and under particular threat from antibiotic use in livestock have been listed by the World Health Organization (WHO) (67). Specifically, these are Fluoroquinolones, 3^rd^ and 4^th^ generation Cephalosporins, Macrolides and Glycopeptides. Empirical evidence relating to antibiotic usage and the evolution, reversion, spread and management of antibiotic resistance should also contribute to guidance decisions. There is an urgent need for quality surveillance studies capable of generating these data.

## Methods

### Mathematical model

The model tracks the spread of an unspecified bacterial pathogen amongst a human population. The impact of the spread of an antibiotic resistance gene from livestock to humans on the evolution and spread of that resistance gene amongst humans is modelled.

The model records the number of individuals in each of five states: treated individuals infected with a sensitive (*T*) or resistant 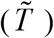 pathogen; untreated individuals infected with a sensitive (*U*) or resistant (*Ũ*) pathogen; and susceptible individuals. Uninfected individuals can become infected with the sensitive or resistant pathogen through exposure to infected humans at a per-person rate equal to the product of the transmission parameter, *β*, and the fraction of individuals infected with each strain. In addition, uninfected individuals can become infected directly with an animal pathogen at a per-person rate of Λ years^-1^. A fraction *α* of these infections are resistant. By varying the relative sizes of *β* and Λ, the model can be parametrized to consider pathogens with different transmission dynamics; for example, those which are primarily transmitted to humans from animals (high *β* and low Λ) and those which are primarily transmitted to humans from other humans (low *β* and high Λ).

A fraction (*π*) of humans receive antibiotic treatment immediately upon infection. Treatment in individuals infected with the drug-sensitive pathogen drives the evolution of the drug-resistant pathogen through *de novo* mutation at a per-person rate of *μ* years^-1^. In addition, the resistance gene can be acquired via gene transfer from a similar or different bacterial genera to the pathogen under consideration from livestock or an alternative (e.g. human or environmental) source at a per-person rate of *ϕ*_*A*_ years^-1^ and *ϕ*_*N*_ years^-1^, respectively. In untreated individuals infected with the drug-resistant pathogen, evolution drives reversion back to the drug-sensitive pathogen at a per-person rate of *Ψ* years^-1^. There are no mixed infections meaning that within-host overgrowth of one strain by another is assumed to occur quickly (68). Recovery of untreated infections occurs at a per-person rate of *δ* years^-1^. Amongst treated individuals, the rate of recovery of antibiotic sensitive (*σ* years^-1^) and antibiotic resistant infections (*ε* years^-1^) are free to vary. Thus, a fitness advantage associated with antibiotic resistance in treated individuals can be modelled through differential recovery rates (by making *ε* < *σ*).

We model the impact of an intervention (intervention 1) in which the prevalence of resistance in livestock is reduced by a proportion *z*_1_, but the total rate that humans are infected by animals remains unchained, thus *α* → *α*(1− *z*_1_). An equivalent reduction in the rate of gene transfer from animals is also assumed, thus *ϕ*_*A*_ → *ϕ*_*A*_ (1− *z*_1_). In Fig S3 and Fig S4 we also investigate the impact of a second intervention (intervention 2) in which we assume that the prevalence of resistance in livestock remains unchanged, but the rate that humans are infected by livestock is reduced by a proportion *z*_2_. Thus Λ → Λ(1− *z*_2_). This could represent a reduction in bacterial infections due to overall improvements in the management of bacterial infections at the farm level or an increase in bacterial testing during food processing. For this intervention the rate of gene transfer from animals is also assumed to be reduced by a proportion *z*_2_, thus *ϕ*_*A*_ → *ϕ*_*A*_ (1− *z*_2_) .

The model is formulated as a set of coupled ordinary differential equations (see equations 1-5) which describe the change over time in the number of individuals in each of the five states. A schematic diagram of the model is shown in Fig 2 and the variables and parameters of the model are listed in Tables 1. The model was simulated using the ode solver ‘ode45’ in the software package Matlab.

### Model Equations in the absence of any intervention

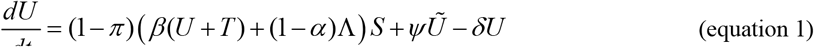

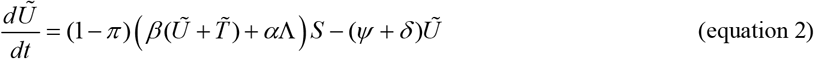

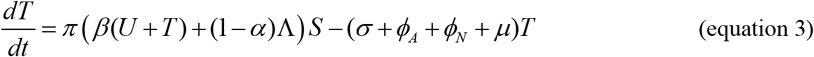

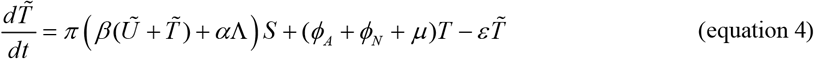

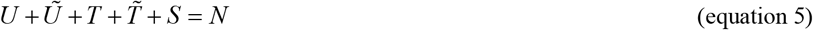

**With intervention 1**: *α* → *α*(1− *z*_1_) and *ϕ*_*A*_ → *ϕ*_*A*_ (1− *z*_1_)

**With intervention 2**: Λ → Λ(1− *z*_2_) and *ϕ*_*A*_ → *ϕ*_*A*_ (1− *z*_2_)

An adapted version of this model in which individuals infected with a resistant strain are further categorized according to whether resistance in the strain originated from livestock or from an alternative (e.g. human) source, is provided in Fig S2. This version is used to evaluate how the impact of interventions targeted at livestock is dependent upon the observed proportion of resistant strains in a particular pathogen that originates from livestock.

## Supporting information captions

**Table S1.** Parameter values used to generate Fig 4. and Fig S3. In addition to the values listed above, for all simulations: *π* = 0.3, *σ* = 24 years^-1^ and *δ* =12 years^-1^. The outcome is independent of *N (*i.e. choose any *N)*. The values of *ϕ*_*A*_, (where non-zero) represent the value prior to either intervention 1 or intervention 2 (i.e. *z*_1_ = *z*_2_ = 0).

| Is there a net fitness cost or benefit to resistance? | What % of observed resistance derives from livestock? | How is the pathogen transmitted to humans? |  |  |
| --- | --- | --- | --- | --- |
| | | Majority from livestock<br>$\beta = 0.0284/N$<br>$\phi_A = 0 \text{ yrs}^{-1}$<br>$\Lambda = 0.1416 \text{ yrs}^{-1}$ | Majority from humans<br>$\phi_A = 0 \text{ yrs}^{-1}$ | All from humans<br>$\beta = 14.26/N$<br>$\Lambda = 0 \text{ yrs}^{-1}$ |
| **Net cost**<br>$\psi = 0.1 \text{ years}^{-1}$<br>$\varepsilon = 23.9 \text{ yrs}^{-1}$ | 5 | $\alpha = 0.0030 \text{ yrs}^{-1}$<br>$\mu + \phi_N = 29 \text{ yrs}^{-1}$ | $\beta = 14.26/N$<br>$\Lambda = 1.11 \times 10^{-4} \text{ yrs}^{-1}$<br>$\alpha = 0.0216$<br>$\mu + \phi_N = 0.138 \text{ yrs}^{-1}$ | $\phi_A = 0.0064 \text{ yrs}^{-1}$<br>$\mu + \phi_N = 0.122 \text{ yrs}^{-1}$ |
| | 95 | $\alpha = 0.0578 \text{ yrs}^{-1}$<br>$\mu + \phi_N = 0.7778 \text{ yrs}^{-1}$ | $\beta = 14.26/N$<br>$\Lambda = 1.11 \times 10^{-4} \text{ yrs}^{-1}$<br>$\alpha = 0.410$<br>$\mu + \phi_N = 0.0073 \text{ yrs}^{-1}$ | $\phi_A = 0.122 \text{ yrs}^{-1}$<br>$\mu + \phi_N = 0.0064 \text{ yrs}^{-1}$ |
| **Neutral**<br>$\psi = 0.0001 \text{ yrs}^{-1}$<br>$\varepsilon = 23.9 \text{ yrs}^{-1}$ | 5 | $\alpha = 0.0030$<br>$\mu + \phi_N = 29 \text{ yrs}^{-1}$ | $\beta = 14.26/N$<br>$\Lambda = 9.98 \times 10^{-5} \text{ yrs}^{-1}$<br>$\alpha = 0.0132$<br>$\mu + \phi_N = 0.0730 \text{ yrs}^{-1}$ | $\phi_A = 0.0034 \text{ yrs}^{-1}$<br>$\mu + \phi_N = 0.064 \text{ yrs}^{-1}$ |
| | 95 | $\alpha = 0.0578$<br>$\mu + \phi_N = 0.7778 \text{ yrs}^{-1}$ | $\beta = 14.26/N$<br>$\Lambda = 9.98 \times 10^{-5} \text{ yrs}^{-1}$<br>$\alpha = 0.255$<br>$\mu + \phi_N = 0.0037 \text{ yrs}^{-1}$ | $\phi_A = 0.064 \text{ yrs}^{-1}$<br>$\mu + \phi_N = 0.0034 \text{ yrs}^{-1}$ |
| **Net advantage**<br>$\psi = 0.0001 \text{ yrs}^{-1}$<br>$\varepsilon = 20 \text{ yrs}^{-1}$ | 5 | $\alpha = 0.003$<br>$\mu + \phi_N = 20.4 \text{ yrs}^{-1}$ | $\beta = 14.20/N$<br>$\Lambda = 3.55 \times 10^{-4} \text{ yrs}^{-1}$<br>$\alpha = 5.43 \times 10^{-5} \text{ yrs}^{-1}$<br>$\mu + \phi_N = 7.30 \times 10^{-5} \text{ yrs}^{-1}$ | $\phi_A = 2.17 \times 10^{-5} \text{ yrs}^{-1}$<br>$\mu + \phi_N = 4.13 \times 10^{-4} \text{ yrs}^{-1}$ |
| | 95 | $\alpha = 0.0558$<br>$\mu + \phi_N = 0.64 \text{ yrs}^{-1}$ | $\beta = 14.20/N$<br>$\Lambda = 3.55 \times 10^{-4} \text{ yrs}^{-1}$<br>$\alpha = 0.001$<br>$\mu + \phi_N = 4.04 \times 10^{-5} \text{ yrs}^{-1}$ | $\phi_A = 4.13 \times 10^{-4} \text{ yrs}^{-1}$<br>$\mu + \phi_N = 2.17 \times 10^{-5} \text{ yrs}^{-1}$ |

**Text S1.**
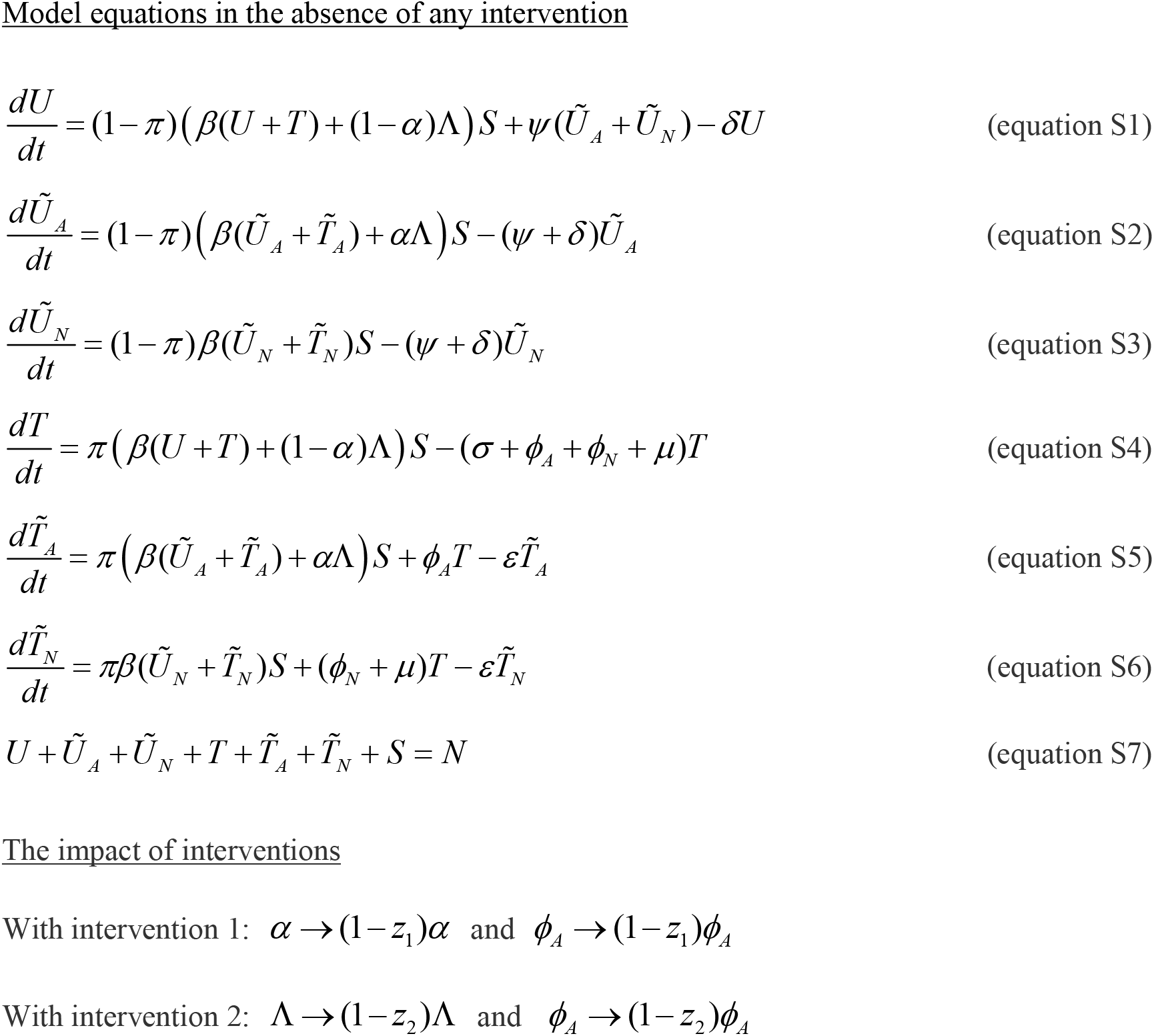
Equations describing a model of antibiotic resistance gene flow from livestock and its spread amongst humans that tracks the origin of resistance.

**Fig S1.**
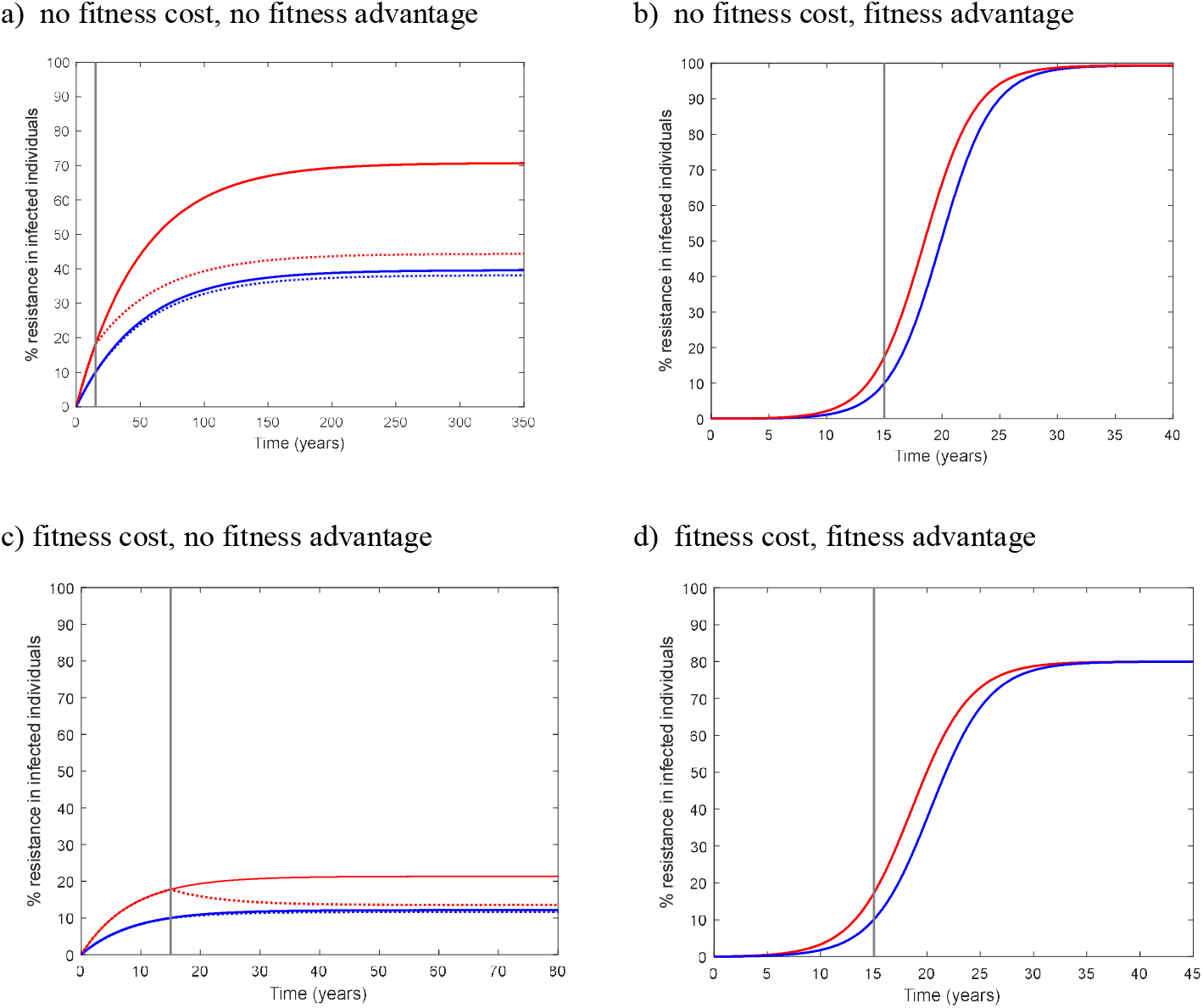
Model predictions of the impact of the spread of antibiotic resistance from livestock to humans on the spread of resistance amongst a primarily human-human spread pathogen. This figure shows how the spread of antibiotic resistance from livestock to humans can contribute to the spread of resistance among humans. It also highlights the expected impact of an intervention (intervention 1) to reduce the spread of resistance from livestock to humans upon resistance prevalence amongst humans. Across panels a)-d) different scenarios in regards to the fitness profile of the resistant strain are compared. The fitness cost of resistance in the absence of treatment is compared by varying the reversion rate. In a) and b) there is a no fitness cost (*Ψ* = 0 years^-1^) and in c) and d) there is a measurable fitness cost (*Ψ* = 0.1 years^-1^) in untreated individuals. The fitness advantage of resistance in the presence of treatment is compared by varying the recovery rate of treated resistant infections. In a) and c) there is no fitness advantage (*ε* = 24 years^-1^), and in b) and d) there is a measurable fitness advantage (*ε* = 20 years^-1^), in treated individuals. In each panel, two rates at which the resistance gene spreads from livestock to humans are compared (solid lines). In blue, most resistance derives from an alternative (i.e. human) source. In red the prevalence of resistance amongst livestock is increased by an amount such that livestock and alternative sources contribute equally to resistance amongst humans. The rates of *ϕ*_*N*_ + *μ* and α differs between a, b, c and d to yield a resistance prevalence of 10% at year 15 when 5% of resistance derives from livestock (blue line). In each figure, at year 15 (grey line), an intervention (intervention 1) that reduces the prevalence of resistance in livestock by 80% (*z*_1_ = 0.8) is applied (see dashed lines where not hidden behind the exact solid lines). In a) and c) the resistant strain does not reach fixation. Reducing the prevalence of resistance in livestock can significantly reduce the prevalence of resistance in humans provided that resistance in livestock contributes measurably to the generation of resistance in humans (compare solid and dashed red lines). In b) eventual fixation of the resistant strain is inevitable. Reducing the rate that resistance is acquired from livestock has no impact upon the prevalence of existing resistant strains. In d) the resistant strain does not reach fixation. Reducing the rate that resistance is acquired from livestock has no measurable impact upon the prevalence of existing strains. The following parameters are used in each simulation: *β* = 14.26, *π* = 0.3, *σ* = 24 years^-1^, *δ* =12 years^-1^, *ϕ A* = 0 years^-1^ and Λ =1.425×^−4^ years^-1^. Furthermore, in a), b), c) and d) respectively, *μ* +*ϕ N* = 0.076, 4.827 ×10^−4^, 0.1383 and 0.0015 years^-1^; in blue*α* = 0.0293, 2.236 ×10^−4^, 0.0527 and 6.8×10^−4^ years^-1^; and in red *α* = 0.53, 0.00425, 0.935 and 0.0130 years^-1^. The outcome is independent of *N* (i.e. choose any *N*).

**Fig S2.**
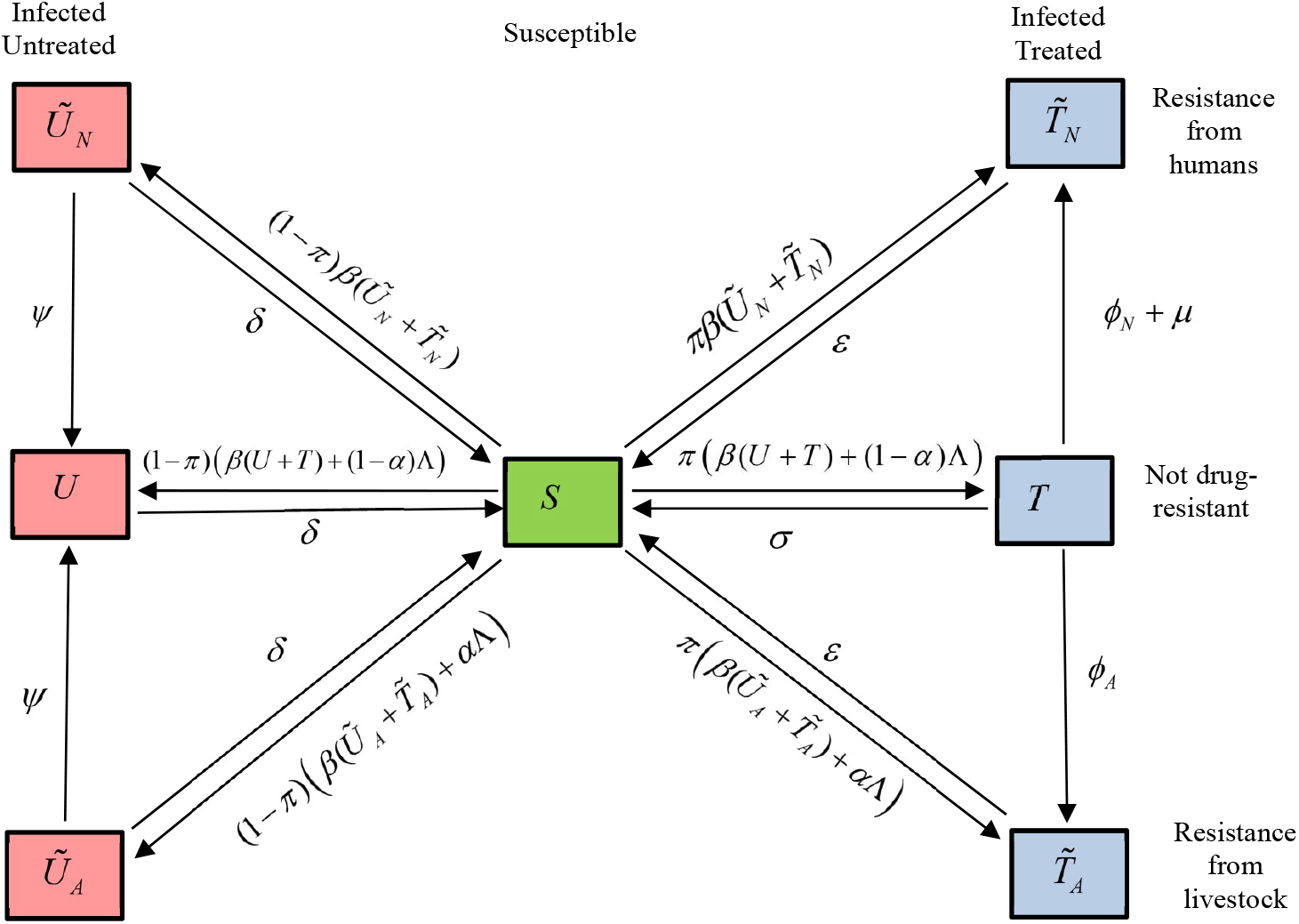
Model 2. A mathematical model of antibiotic resistance gene flow from livestock and its spread amongst humans that tracks the origin of resistance. This model tracks the spread of a bacterial pathogen amongst a human population, a portion of which receive antibiotic treatment upon infection. Infections can result from transmission from other infected humans or from exposure to infections in livestock. There is competition between antibiotic-sensitive and antibiotic-resistant strains. Resistant strains can emerge in treated individuals via *de novo* mutation or gene transfer deriving from livestock or an alternative (i.e. human) source. In addition, resistant infections can be transmitted to humans from livestock directly. Resistant strains can revert in untreated individuals. Recovery of infection occurs at different rates according to the pathogen strain and the treatment status. This model is an expanded version of the Model 1, presented in the main text. It additionally records whether resistance derives from livestock or an alternative source. 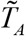 and *Ũ*_*A*_ record the number of treated and untreated humans infected with a resistant strain that derives from livestock, respectively. 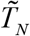 and *Ũ*_*N*_ record the number of treated and untreated humans infected with a resistant strain that derives from an alternative source, respectively. The remaining variables and parameters of the model are the same as in Model 1. The impact of interventions to reduce the spread of resistance from livestock to humans is excluded from this illustration but is included in the model in the same way as for Model 1 (see Methods). Model equations are provided in Text S1.

**Fig S3.**
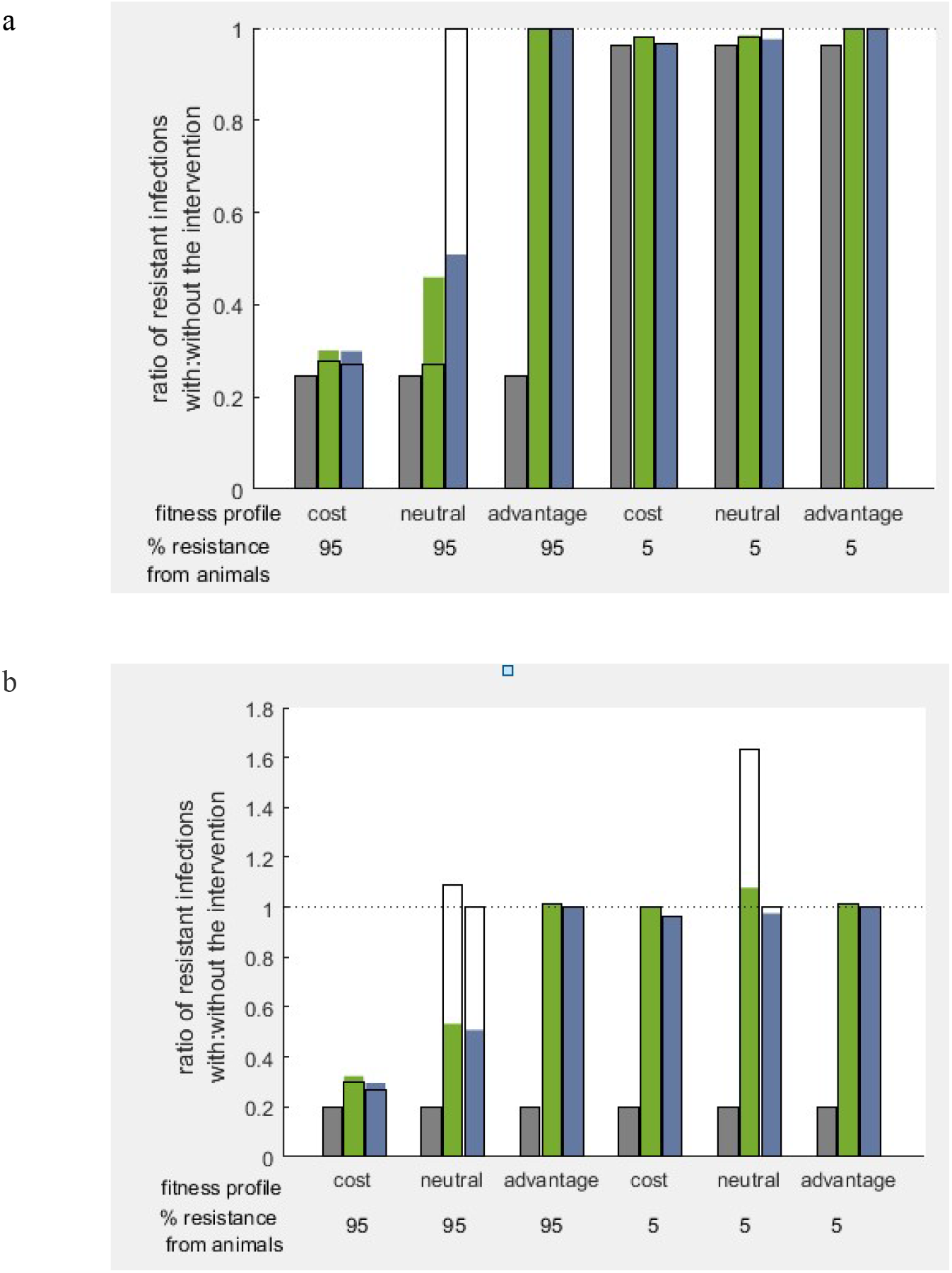
Whether the number of human infections with an existing antibiotic resistance strain can be reduced by interventions aimed at livestock depends upon different factors. This figure explores the impact that livestock-targeted interventions have on the number of human infections with existing resistant strains, according to different factors. This figure is an extension of figure 4, but it also considers the following additional considerations: the impact of a second intervention (intervention 2); strains with a neutral fitness cost associated with resistance; and the long term impact, i.e. at equilibrium. The factors that are explored in this figure are the transmission dynamics of the pathogen, the fitness profile of the resistance strain and the percentage of observed resistance that derives from livestock. The coloured bars categorize whether the pathogen is transmitted to humans primarily (99.9%) from animals (grey), primarily (99.9%) from humans (green) or entirely from humans (blue). The fitness profile is varied to consider a resistant strain that either has a net fitness cost to resistance (measurable fitness cost in untreated individuals and negligible fitness advantage in treated individuals), neutral fitness (negligible fitness cost, negligible fitness advantage) or a net fitness advantage to resistance (negligible fitness cost, measurable fitness advantage). In each model simulation used to generate the results shown, the model is independently parametrized so that prior to time 0, an infection has reached a prevalence of 1% amongst humans and antibiotic resistance is absent. At time 0, antibiotic resistance emerges amongst humans at such a rate that 15 years later, 10% of infections are resistant. In one option, 95% of resistance observed at year 15 originates from livestock, whereas in the other option the value is 5%. Intervention 1 (panel a) reduces the prevalence of resistance amongst zoonotic infections by 80%. Intervention 2 reduces the rate of zoonotic infections by 80%. Both interventions reduce the rate of antibiotic gene transfer from animals to humans by 80%. The coloured bars show the ratio of the number of resistant human infections at year 50 with:without the intervention. The black outlines of bars show the same ratio calculated at equilibrium. The full set of parameters used for these simulations are listed in Table S1.

**Fig S4.**
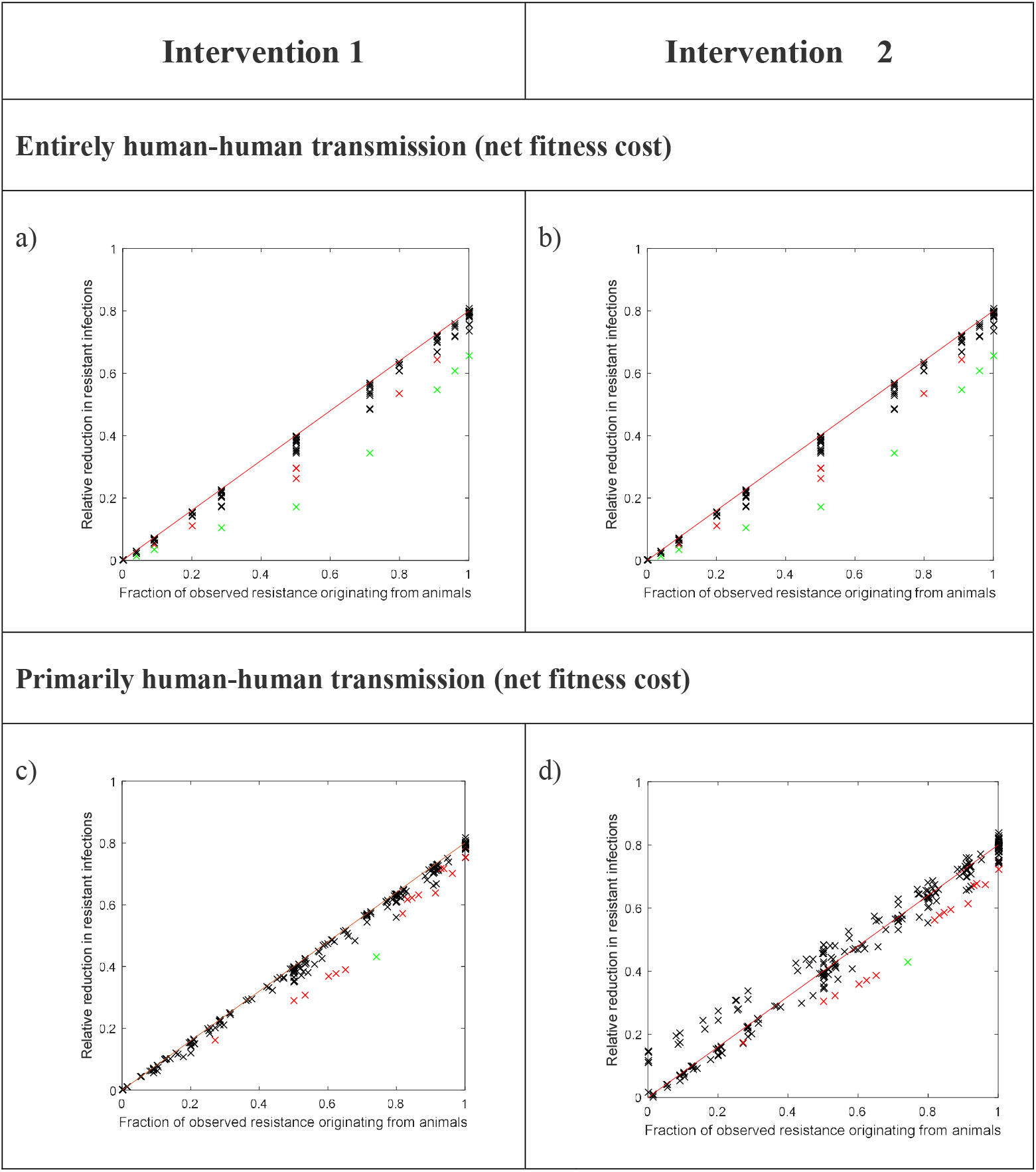
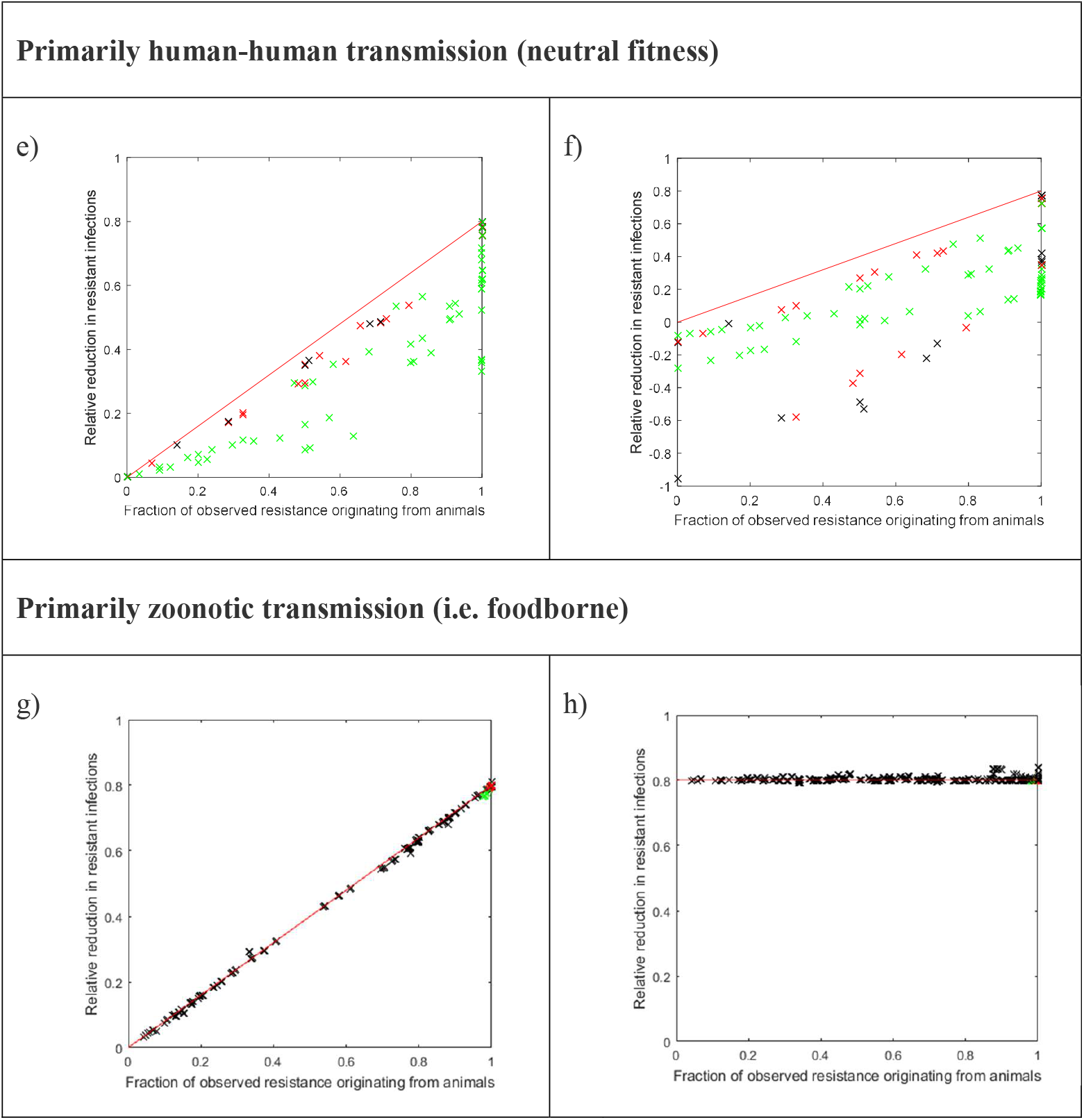
The fraction of observed resistance that originates from livestock is an important determinant of the relative reduction in the number of human infections with existing resistance strains that could be achieved through intervenetions targeted at livestock. The model was simulated under a broad range of parameter combinations to understand how the reduction in the equilibrium number of resistant human infections resulting from intervention 1 (panels a, c, e and g) or intervention 2 (panels b, d, f and h) is dependent upon the equilibrium prevalence of resistance in that pathogen that derives from livestock. In each simulation, we assume that human exposure to resistance in livestock is reduced by 80% (*z*_1_ = 0.8 for strategy 1 and *z*_2_ = 0.8 for strategy 2). Panels a-d concern pathogens that entirely (a and b) or primarily (c and d) transmit to humans from humans and for which there is a net fitness cost to resistance. In these scenarios, provided the equilibrium prevalence is low, (<30% black markers) the number of resistant infections is approximately reduced by a proportion *z*_1_ *p* for strategy 1, or *z*_2_ *p* for strategy 2 (shown by the red lines). In this expression, *p* represents the fraction of resistance at equilibrium that derives from livestock. For larger equilibrium prevalences (30-50% red markers and >50% green markers), the reduction can smaller than *z*_1_ *p* or *z*_2_ *p*, respectively. Panels e-f concern pathogens that primarily transmit to humans from humans and for which resistance has neutral fitness. In these scenarios, Intervention 1 can reduce the equilibrium number of resistant infections amongst humans. Intervention 2 can change the equilibrium number of resistant infections, but not necessarily reduce it. Panels g-h concern pathogens that primarily transmit to humans from animals (i.e. foodborne pathogens). In these scenarios, the number of resistant infections is approximately reduced by a proportion *z*_1_ *p* for strategy 1 or *z*_2_ for strategy 2 (shown by the red lines).

